# Modelling PRRS transmission between pig herds in Denmark and prediction of interventions impact

**DOI:** 10.1101/2025.05.09.653077

**Authors:** You Chang, Ana Rita Pinherio Marques, Mette Fertner, Nils Toft, Bjørn Lorenzen, Mossa Merhi Reimert, Hans Houe, Beate Conrady

## Abstract

Porcine reproductive and respiratory syndrome (PRRS) is an endemic viral disease in most pig-producing countries, including Denmark. In 2022, Denmark launched a control program to reduce PRRS prevalence, with legislative changes in 2023 making testing and status reporting mandatory. The program also enforces the loss of PRRS-free status for farms that purchase pigs from non-PRRS-free sources and implements region-specific control measures to coordinate PRRS elimination within herds. However, the effectiveness of these interventions remains uncertain and requires thorough evaluation through transmission modelling and analysis of data before and after legislation changes.

To understand PRRS transmission prior to legislative changes in 2023 and predict the impact of control measures, we developed a between-herd stochastic compartmental model. This model includes compartments for susceptible (*S*), highly infectious (*I*_*h*_), lowly infectious (*I*_*l*_) and detected (*D*) pig herds, using data from 2020-2021. The model (i) quantifies the relative contributions of pig movements and local transmission to the spread of PRRS; (ii) generate herd-level maps of the basic reproduction (*R*_0_); and (iii) assess the effectiveness of targeted interventions for eradicating PRRS in Denmark.

Model results indicated that more than 50% of herds had an *R*_0_ greater than 1, suggesting a potential for sustained transmission if no interventions had been implemented after 2022. Both local spread and movement-mediated transmission play important roles, but local transmission drives the spatial heterogeneity in observed PRRS prevalence across Denmark. Although only 17% of infectious herds remain undetected under current surveillance, they are responsible for 60% of total transmission. Local control via depopulation and repopulation, is the fastest measure to reduce the observed prevalence of PRRS, but it has a lower effect on true transmission due to the hidden infections. Therefore, achieving eradication may require a combination of more frequent testing, targeted within-herd PRRS elimination and stricter risk-based trading. This study identifies PRRS hotspots and transmission routes, offering evidence-based recommendations for control.

**Highlights:**

- We generated the first between-herd *R*_0_ map of PRRS transmission in Denmark
- 17% of undetected herds are responsible for 60% of total PRRS transmission
- Positive stable herds are 50 times less infectious than positive unstable herds
- Local spread drives the spatial heterogeneity of PRRS transmission
- Eradication requires multiple controls e.g., local controls, stricter risk-based trading and more frequent testing

## 1. Introduction

Porcine reproductive and respiratory syndrome (PRRS) is a viral disease that affects pig populations worldwide (Lunney et al., 2010). The disease causes significant production losses due to reproductive failure in sows, respiratory distress in younger pigs, and increased mortality rates (OIE, 2008). PRRS is estimated to cause production losses ranging between €4 to €139 per sow per year in Denmark (Rathkjen & Dall, 2017), €126 per sow in the Netherlands (Nieuwenhuis et al., 2012) and $664 million in losses annually for the pig industry in the U.S. (Holtkamp et al., 2013). The infection spreads through multiple routes, including animal movement, indirect contacts such as contaminated materials, and airborne transmission (Cho & Dee, 2006; Corzo et al., 2010; Rahe & Murtaugh, 2017; Rowland & Morrison, 2012; Valdes-Donoso et al., 2018). The etiological agent of PRRS is an RNA virus, the porcine reproductive and respiratory syndrome virus (PRRSV), which is divided into type 1 and type 2 genotypes, known as European and North American strains, respectively (Murtaugh et al., 1995; Rahe & Murtaugh, 2017).

Denmark has faced endemic PRRS transmission since its first detection in March 1992 (Bøtner et al., 1994). The infections were widespread by mid-1990s, with a herd-level prevalence of approximately 25% in sow herds and 33% in finisher herds, with an annual herd-level incidence of 5-10% (Mortensen, 1996; Mortensen et al., 2002; Mortensen & Strandbygaard, 1996). Denmark’s PRRS control efforts have evolved over time, with the Specific Pathogen Free (SPF) system forming the foundation of PRRS management since 1993 (Fertner et al., 2025). The SPF system is an industry-driven voluntary system. Enrolled herds are routinely tested for the absence of seven pathogens, including PRRS, as part of the health declaration system. To sustain the SPF status of herds, herds must undergo annual biosecurity evaluations and follow a certain set of rules regarding biosecurity (Lopes Antunes et al., 2019; Conrady et al., 2023; SPF, 2024), including investments in changing farm facilities, routine serological testing, herd status classification, and restrictions on animal movement. Despite of these efforts, national PRRS prevalence remained around 35% in 2020 (Kristensen et al., 2020; Sundhedsstyringen, 2020).

In May 2022, Denmark launched national PRRS control program that extended PRRS control beyond the SPF program. A key legislative change came in 2023 when a government order was issued, obligating all herds to officially declare their PRRS status (Danish Ministry of Food, Agriculture and Fisheries, 2023; State Serum Institut, 2023). In addition, local control measures have been introduced, encouraging herd owners to collaborate and coordinate actions to eliminate PRRS in positive herds within specific local areas. The final decision on whether, and when, to eliminate PRRS within a herd lies with the herd owners. These decisions can be influenced by within-herd outbreak size and economic considerations, rather than a compulsory standard control. While rolling out these initiatives, the effectiveness of interventions after new legislation remains uncertain.

To predict the effectiveness of interventions targeting different transmission routes, it is essential to understand the relative contributions of each route, including pig movements, and local spread. Local spread occurs when virus is transmitted through contact with virus-laden fomites (such as shared equipment, vehicles, and workers) or virus-laden aerosols in the vicinity of an infectious herd, often referred to as airborne transmission. Several mechanistic modeling studies in the U.S. (Galvis, Corzo, & Machado, 2022; Galvis, Corzo, Prada, et al., 2022) and Canada (Thakur, Revie, et al., 2015; Thakur, Sanchez, et al., 2015) have highlighted the importance of these routes in the between-herd spread. Herd-level risk factor analyses have identified regional swine density and the purchase of PRRSV-positive animals as primary risk factors (Baysinger et al., 1997; Firkins & Weigel, 2004; Mortensen et al., 2002; Velasova et al., 2012). However, these findings cannot be directly applied to other regions or situations where interventions alter disease dynamics. The relative contribution of local spread and movement may vary between countries due to demographic and environmental heterogeneity, such as livestock density, frequencies of livestock movements, herd size, farm biosecurity standards, and farm structures (Conrady et al., 2023, 2024). Therefore, a mechanistic model specific to Denmark is needed to understand transmission and predict the impact of new legislative changes (after 2023) on PRRS control.

The present study aims to develop a stochastic compartmental model to *(i*) generate herd-level maps of the basic reproduction ratio (*R*_0_); (*ii*) quantify the relative contributions of pig movements and local transmission to the spread of PRRS; and (*iii*) assess the effectiveness of targeted interventions for PRRS control in Denmark.

## 2. Material and method

We developed a stochastic between-herd compartmental model to represent the PRRS transmission dynamics using the SimInf package (Widgren et al., 2016; Widgren et al., 2019). The data were described in Section 2.1, followed by the model description in Section 2.2. In addition, parameters were explained in Section 2.3, and statistical method was described in Section 2.4, and the calculation of the basic reproduction ratio (R_0_) in Section 2.5. In Section 2.6, uncertainty in the parameters was investigated by sensitivity analysis, and in Section 2.7, several interventions that target different transmission routes were explained.

### 2.1. Data

Data used in this study covered the period from January 1, 2020, to December 31, 2021, and consisted of 4 main datasets: (1) herd-level PRRS status and biosecurity level extracted from the Danish SPF health dataset; (2) production type; (3) the number of moved pigs between premises considered as discrete herd-to-herd pairs and (4) geographic information of the herd locations (Figure 1).

**Figure 1.**
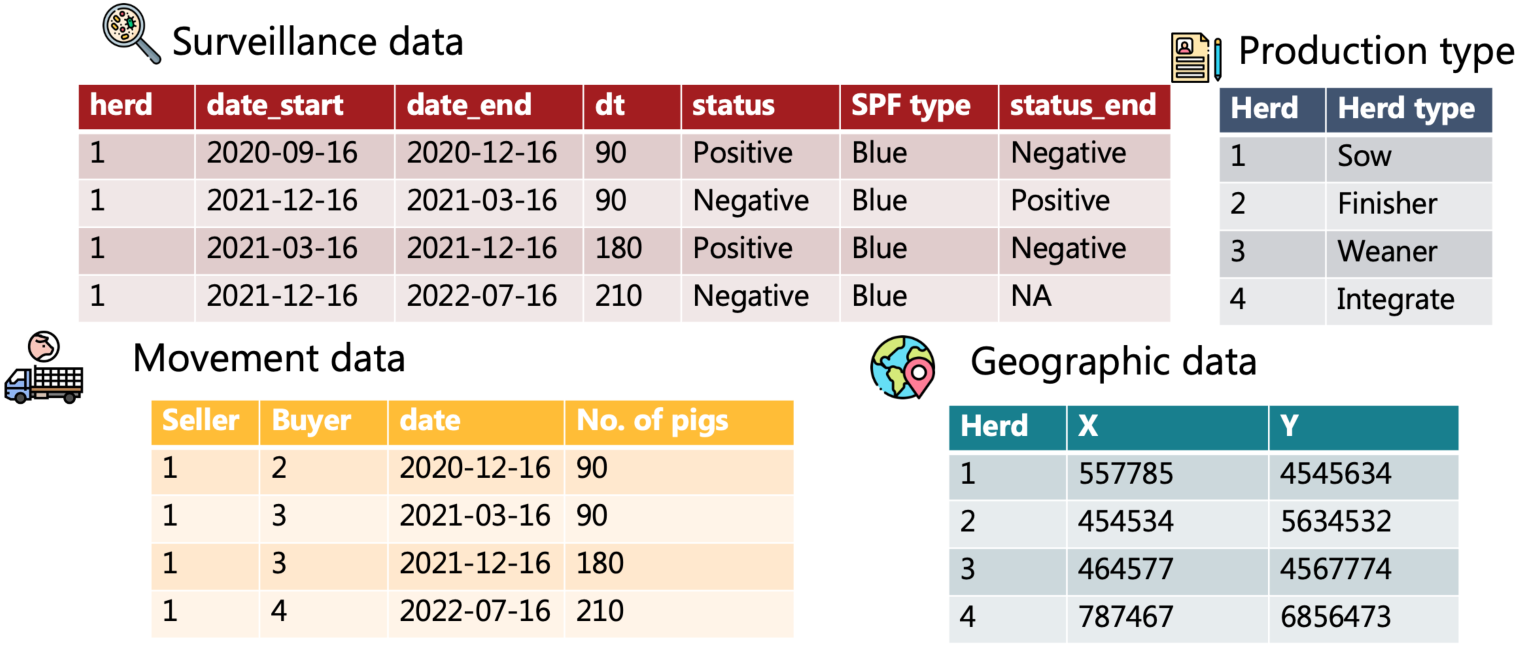
Example of four synthetic datasets, illustrating data structures used in this study.

The study population consists of herds with reported PRRS status during the study period (2021-2022). In total, 3,420 herds were included, representing approximately 60% of all pig herds in Denmark during this period. For each herd, we extract the associated herd identification number, biosecurity level, and the PRRS status.

In SPF system, herds’ biosecurity level is classified into green, blue and red level and non-SPF herds (Lopes Antunes et al., 2015):

1. Red herds: These herds undergo monthly testing and demonstrate a high commitment to eliminating PRRS in the event of an outbreak.
2. Blue and green herds: These herds undergo annual testing and show a moderate commitment to achieving PRRS elimination.
3. Non-SPF herds: These herds have the lowest biosecurity measures and typically do not report their status. However, some non-SPF herds had data entries in the surveillance system, which were also included in this study.

For each testing, samples from 10 animals (5 gilts and 5 sows) from red herds and 20 animals from blue herds (selected randomly within the herd) are collected and tested by serological tests. The current SPF health system automatically assigns a PRRS-positive status to herds receiving pigs from PRRS-positive herds without additional testing. While this system is designed to reduce risk, it increases the proportion of false positive test results in the official control system.

### 2.2 Model description

A stochastic between-herd compartmental model was developed using R package SimInf (Widgren et al., 2016; Widgren et al., 2019). In this model, herds are considered as epidemiological units and are classified into four compartments (Figure 2): Susceptible (*S*), highly infectious (*I*_*h*_), lowly infectious (*I*_*l*_) and detected (*D*). When a susceptible herd is introduced to PRRSV, it typically enters an acute epidemic phase, during which most pigs become infected in a short period. This phase is referred to as the highly infectious herds, characterized by significant viral shedding and a high risk of infecting other herds. This corresponds to “positive unstable” herds in PRRS terminology (Holtkamp et al., 2011). Over time, typically after approximately three months (expert’s opinion), the infection stabilizes and becomes endemic within the herd, but herds remain seropositive. Herds in this stable phase, referred to as the lowly infectious herds, have much lower levels of viral shedding and a lower risk of infecting other herds. This corresponds to “positive stable” herds in PRRS terminology (Holtkamp et al., 2011). The transition from the highly infectious phase to the lowly infectious phase is modelled as *γI*_*h*_. Both infectious phases can contribute to the local spread and movement-mediated transmission.

**Figure 2.**
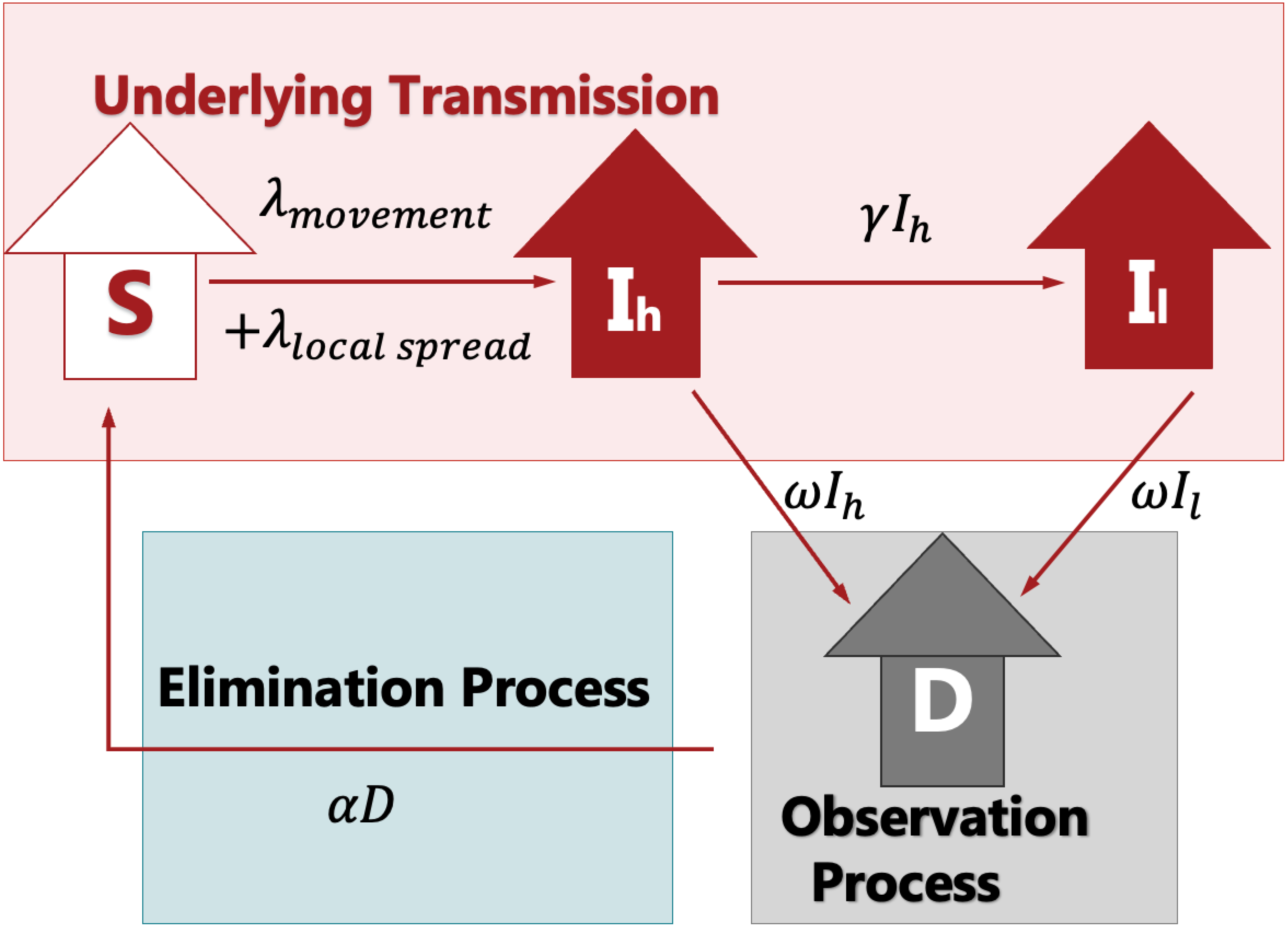
A conceptual diagram of between-herd PRRS transmission. Transmission occurs through local spread and movement. Local spread occurs within a 5 km radius due to indirect contact and airborne transmission at a rate of *λ*_*local* *spread*_. Movement-mediated transmission, modelled using trading data, occurs at a rate of *λ*_*movement*_. Highly infectious herds can transition to lowly infectious (seropositive) herds at a rate of *γI*_*h*_. Both highly and lowly infectious herds can be detected as positive through blood testing and clinical checkups. Upon detection, herds can implement within-herd PRRS elimination at rate of *αD*.

The transmission rate via movement (*λ*_*movement*_) is modelled as *P*_*mov*_ ⋅ *I*_*h*_*mov*__ + *P*_*mov*_ ⋅ *k*_*mov*_ ⋅ (*I*_*lmov*_ + *D*_*mov*_/,

where:

- *P*_*mov*_represents the infection probability per risky movement from a highly infectious herd
- *I*_*hmov*_ represents the number of risky movement contacts with the highly infectious herds per day
- *k*_*mov*_ is a scaling factor representing the relative infectiousness of lowly infectious compared to highly infectious herd via movement-mediated transmission
- *I*_*lmov*_ represents the number of movement contacts with lowly infectious herds
- *D*_*mov*_ represents number of movement contacts with detected herds.

Under the SPF system, risk-based trading is encouraged and practiced on a voluntary way. Herds with a higher biosecurity level (i.e. red herds) can sell pigs to those with a lower biosecurity level (i.e. blue herds), but not vice versa (Supplement Figure S1). Additionally, herds lose their PRRS-negative status if they acquire pigs from PRRS-positive herds. This means that risky movements - buying pigs from positive herds-can still occur, as herd owners decide whether to find a new trade partner with PRRS-negative status or continue to trade while accepting the loss of their PRRS-negative status. Therefore, in the default scenario, representing the period from 2020 to 2021, we model a proportion of herds complying with risk-based trading rule by purchasing pigs only from PRRS-negative herds. The level of compliance is determined using surveillance data.

Transmission via local spread is caused by contact with virus-laden fomite (shared equipment, vehicle, and workers), or virus-laden aerosols in the vicinity of an infectious herd (often referred to as airborne transmission). In this study, a local area is defined as a 5 km radius area, within which the transmission rate is assumed to be density-dependent; hence the transmission rate (*λ*_*local* *spread*_) is modelled as *β*_*ls*_ ⋅ *I*_*h*_*ls*__ + *β*_*ls*_ ⋅ *k*_*ls*_ ⋅ ( *I*_*l_ls_*_ + *D*_*ls*_),

Where:

- *β*_*ls*_ represents the transmission rate parameter for local spread of highly infectious herds
- *I*_*h_ls_*_ represents the number of highly infectious neighbor herds (within 5 km of the recipient herd)
- *k*_*ls*_ is a scaling factor representing the relative infectiousness of lowly infectious compared to highly infectious herd via local spread
- *I*_*l_ls_*_ represents the number of lowly infectious neighbor herds (within 5 km of the recipient herd)
- *D*_*ls*_ represents the number of detected PRRS positive herds (within 5 km of the recipient herd).

Both types of infectious herds can be detected via blood testing (active surveillance) or clinical checks by veterinarians (passive surveillance). Regular blood sample testing is conducted, with the frequency of sampling determined by herds’ biosecurity level. In addition, herds are visited by veterinarians regularly, with the frequency and sensitivity of clinical checkups determined by herds’ production types, as sow herds have more severe clinical signs than finisher and weaner herds. Therefore, the detection of PRRS is modelled as *blood*_*sampling*_ ⋅ *SE*_*b*_ + *clinical*_*visit*_ ⋅ *SE*_*clinic*_, where:

- *blood*_*sampling*_ is the rate of regular blood sampling
- *clinical*_*visit*_ is the rate of the vet visiting for clinical check up
- *SE*_*b*_ and *SE*_*clinic*_ are the sensitivities of blood sampling and clinical checkups.

After detection, PRRS positive herds can eliminate the virus through control measures (see Section 2.7). The time required for elimination depends on when farmers implement control measures and the specific strategies they choose. Once PRRS is eliminated, herds are assumed to become immediately susceptible. The elimination is modelled as *αD*, where *α* represents the elimination rate determined by the SPF types (see Table 2).

### 2.3 Parameters

The model includes two types of parameters: local and regional. Local parameters are herd-specific and reflect characteristics such as biosecurity levels, production type, trading pattern matrix, and proximity to neighboring farms, as well as farm management decisions related to interventions (Table 1).

**Table 1.**
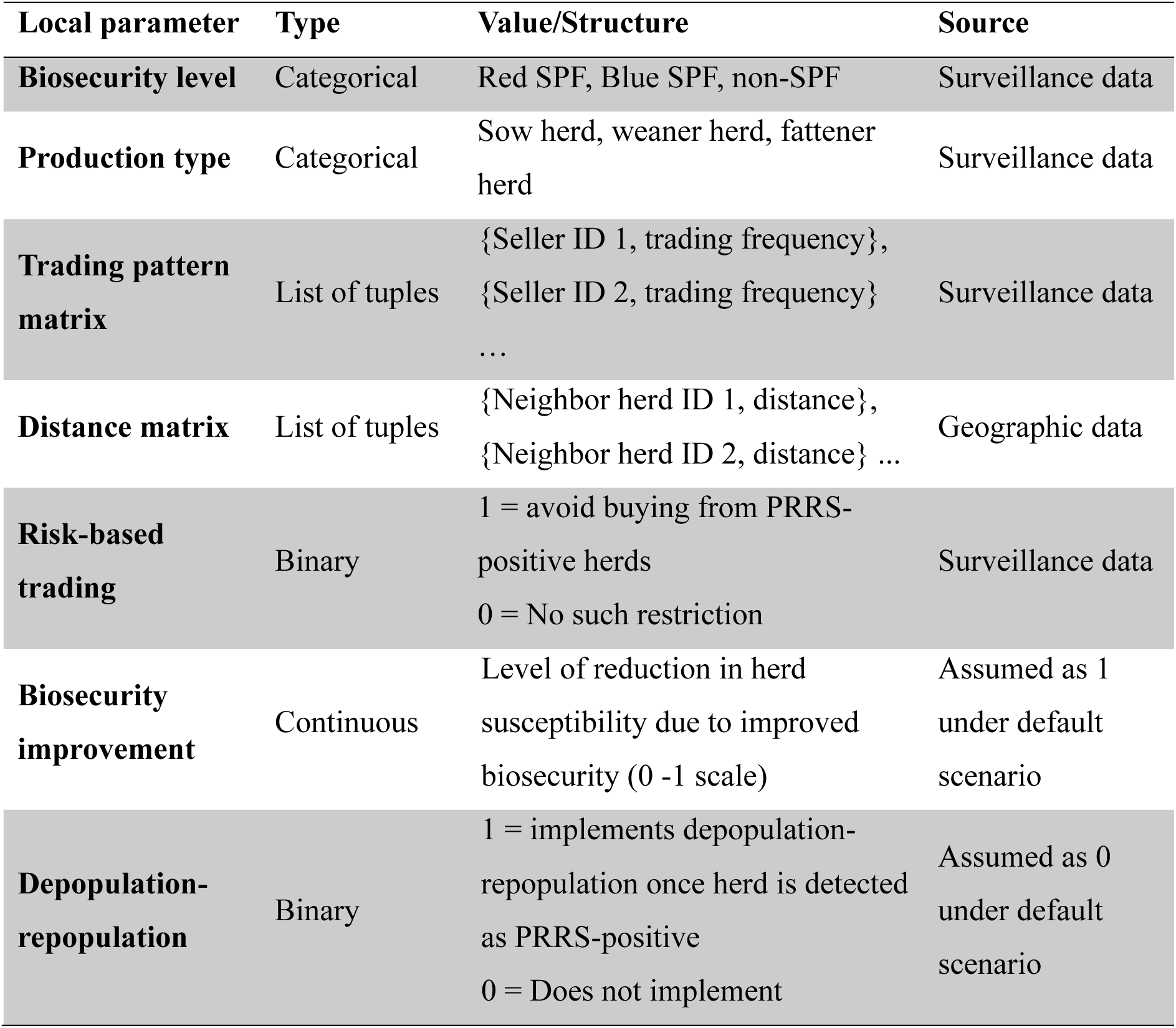
Local parameters of the model.

Regional parameters remain consistent across all areas and include transmission rate parameter for local spread and infection probability via movement, the infectious period of herds, and sensitivity for blood testing and veterinary clinical checks (Table 2). Due to discrepancies between two references regarding to herd-level sensitivity of clinical checkup, the impact of using the alternative herd-sensitivity is analyzed and presented in Supplement Figure S2.

**Table 2.**
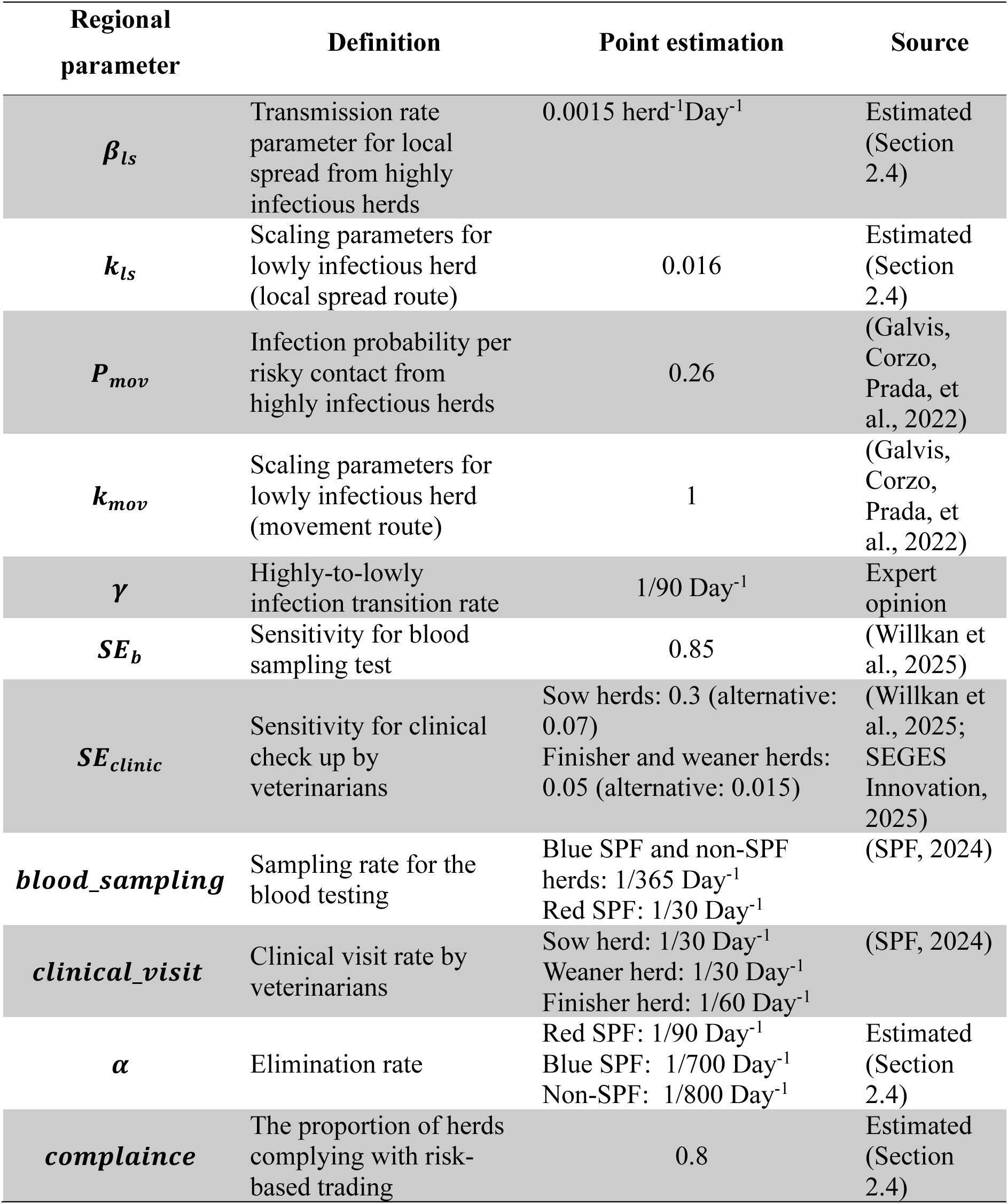
Regional parameters of the model.

### 2.4 Statistical method

We estimate transmission rate parameters for local spread by fitting time-series infection data into the model. This method relates the exposure to the infection sources to the infection probability. The number of herds becoming infected over each test interval (*t*, *t* + *Δt*) follows a binomial distribution with a binomial total susceptible herd at each time interval (which is the time difference between two data entries in surveillance data). The probability of getting infected due to local spread can be derived as:

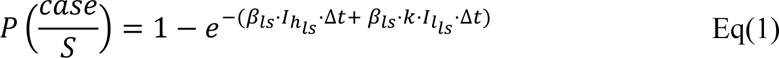

Where 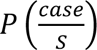 is a binary value representing whether a new case occurs during an observation time interval. *β*_*ls*_ ⋅ *I*_*h_ls_*_ ⋅ Δ*t* represents the transmission rate contributed by highly infectious herds within a 5 km during a time interval Δ*t* while *β*_*ls*_ ⋅ *k* ⋅ *I*_*l_ls_*_ ⋅ Δ*t* accounts for the transmission rate from lowly infectious herds within the same radius and time interval.

The likelihood function, as a function of transmission rate parameters, is given by:

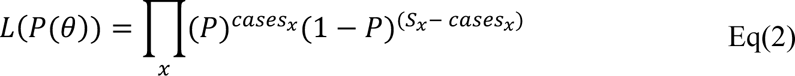

Where P is derived from Eq(1). *θ* represents the two parameters *β*_*ls*_ and *k*_*ls*_. The index *x* donates the number of data lines in the infection dataset. The confidence interval of estimation was listed in the Supplement Table S3.

The elimination rate *α* per SPF types is estimated by calculating the average time between the detected and the return to negative herd. Similarly, the compliance to risk-based trading is calculated as the proportion of herds that did not receive pigs from positive herds.

### 2.5 *R*_0_ calculation

In this study, we calculate two types of *R*_0_, including between-herd *R*_0_ for each herd and a national *R*_0_ (Chang et al., 2025). Between-herd *R*_0_ represents the number of herds that an infectious herd can infect in a population where all other herds are susceptible, which is used to present the spatial heterogeneity. We obtain the between-herd *R*_0_ using Next Generation Matrix method (Diekmann et al., 2010) (Supplement Text S4). The between-herd *R*_0_consists of two components: the local spread route *R*_*local* *spread*_ and movement route *R*_*movement*_:

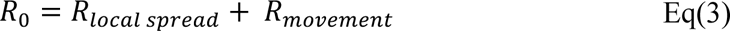

For local spread route, the *R*_*local* *spread*_ can be calculated as:

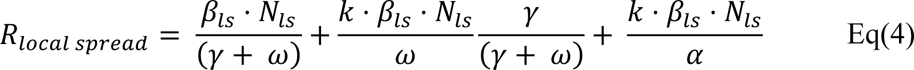

Where the first part of the equation represents the contribution from highly infectious herds while the second part and the third part represent the contribution from lowly infectious herds (undetected or detected part respectively). *N*_*ls*_is the total number of herds in a 5 km neighborhood. *γ* is the rate at which herds transition from highly infectious to less infectious but seropositive phases. *ω* represents the average detection rate, which varies by biosecurity and production type. It is calculated based on the frequency and sensitivity of test or clinical checkups as below:

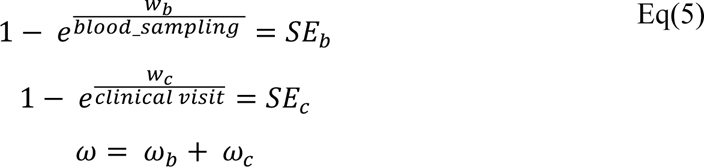

In the Eq (4), the 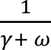 represents the average time herds stay in a highly infectious phase. The 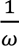 represents the period that herds remain in the lowly infectious phase. The ratio 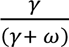 represent the proportion of herds that reach the lowly infectious phase before detection. Similarly, *R*_*movement*_ can be calculated as:

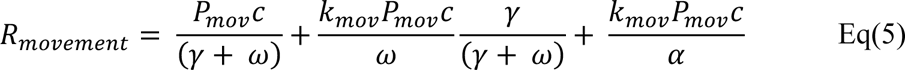

where *c* is contact rate with the outward trading partners.

The national level *R*_0_ represents the average number of herds that one infectious herd can infect when all the other herds are susceptible in Denmark. National level *R*_0_ is calculated using the simulated herd prevalence once transmission dynamics reach equilibrium 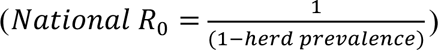 (Chang et al., 2025). For each intervention strategy, a corresponding national *R*_0_can be estimated. If simulated herd prevalence decreases to 0, it indicates that PRRS can be eradicated by the intervention (*R*_0_ < 1). However, if herd prevalence decreases but reaches a new equilibrium, the intervention can reduce transmission but cannot eradicate PRRS (national *R*_0_ > 1).

### 2.6 Sensitivity analysis

Under the current surveillance system, herds lose their PRRS-negative status when they acquire pigs from PRRS-positive herds, assuming a 100% infection risk from such movements without testing. This assumption in the surveillance data prevents estimation of actual the infection probability per movement (*P*_*mov*_) via pig movements and surveillance data. To address this, we adopt literature-based estimate for *P*_*mov*_ in our default scenario (Scenario S1). In Scenario S2, we apply expert opinion from the Danish pig industry, setting *P*_*mov*_to 1 and assuming scaling factor *k*_*mov*_is the same as the *k*_*ls*_(Table 4; Scenario S2). In Scenario S3, reflects the current surveillance assumption, with both *P*_*mov*_ and *k*_*mov*_set to 1, while halving the estimated local transmission parameters estimated from Eq. (2). The values used in all three scenarios are summarized in Table 4.

**Table 3.** Detection rate based for herds categorized with different production types and SPF types.

| Parameter | Definition | Value |
| --- | --- | --- |
| $\omega_b$ | The average detection rate for blood sampling | Red SPF: 0.0630 Day<sup>-1</sup> |
|  |  | Blue SPF: 0.0052 Day<sup>-1</sup> |
|  |  | Non-SPF: 0.0052 Day<sup>-1</sup> |
| $\omega_c$ | The average detection rate for clinical checkup | Sow herds: 0.0120 Day<sup>-1</sup> |
|  |  | Weaner herds: 0.0017 Day<sup>-1</sup> |
|  |  | Finisher herds: 0.00085 Day<sup>-1</sup> |

**Table 4.** Transmission rate parameters and infection probability for sensitivity analysis.

| Parameter(unit) | Scenario 1 | Scenario 2 | Scenario 3 |
| --- | --- | --- | --- |
| $P_{mov}$ (unitless) | 0.26 | 1 | 1 |
| $k_{mov}$ (unitless) | 1 | 0.016 | 1 |
| $\beta_{ls}$ (per herd<sup>-1</sup>Day<sup>-1</sup>) | 1.5e-3 | 1.5e-3 | 7.5e-4 |
| $k_{ls}$ (unitless) | 0.016 | 0.016 | 0.016 |

### 2.7 Interventions

Interventions can be classified into several groups, including those targeting internal biosecurity, external biosecurity, movement and testing frequency. Improving internal biosecurity can shorten a herd’s infectious period. For instance, depopulation-repopulation (DPRP) is an effective but costly method of PRRS elimination within herds and is mainly used in herds with weaners and finishers. This approach involves fully depopulating a production unit, thoroughly cleaning and disinfecting the facilities, and then introducing new PRRS-free pigs after seven days of being empty. This strategy can return a sow herd to susceptible status in approximately 5 months, a weaner herd in 2 months, and a finisher herd in 3 months, depending on the production schedule. To model this intervention, each herd has a binary depopulation-repopulation parameter indicating the herd owners’ decision (See Table 1).

In comparison, external biosecurity, such as quarantine facilities, disinfecting trucks and controlled ventilation, can reduce the transmission risk between herds. The SPF system provides a rough classification of farm biosecurity levels. However, since we cannot find significant difference in the transmission rate parameters among different SPF system herds, we assume that the biosecurity level is consistent across herds. Given that we do not have the quantified data on the impact of each biosecurity measure on transmission risk, nor the percentage of farms that comply with each measure, we model the effect of improved external biosecurity on transmission rate parameter by a factor between 0 and 1 (Table 1), and the number of herds adopting improved external biosecurity is modelled to vary from 0% ∼ 100%.

Herds are recommended not to purchase pigs from PRRS-positive herds. Therefore, each herd is assigned a binary local parameter indicating whether it follows this recommendation (Table 1). During the study period, about 80% of herd owners complied with the risk-based trading rule and 20% of herds received pigs from PRRS-positive herds, which is modelled in the default scenario. Lastly, the active and passive surveillance system is modelled with sensitivity and frequency presented in Table 2.

In addition to the default scenario, we investigate several additional interventions that target different routes:

Interventions at movement route:

1. Default scenario: risk-based trading with 80% of compliance
2. Risk-based trading with 100% compliance: Herds only buy pigs from herds that have not been detected as positive. However, they might still acquire pigs from infectious but undetected herds (*I*_*h*_and *I*_*l*_)
3. Movement ban: The scenario is for exploration

Interventions reduce the external biosecurity, hence reducing susceptibility of herds:

1. Default scenario: under the default biosecurity (red and blue)
2. 50% of herds are randomly selected to enhance external biosecurity, reducing their susceptibility to PRRS infection by 20%
3. All the herds in high-risk herds will improve their external biosecurity, reducing the herds susceptibility by 20%

Interventions improve the internal biosecurity via DPRP (i.e. depopulation-repopulation):

1. Implement DPRP in 50% of herds, selected randomly
2. Implement DPRP in the high-density areas (herds with > 10 neighboring herds within 5km)
3. Implement DPRP in all herds

Interventions at surveillance system:

1. Increase testing frequency to every half year
2. Increase testing frequency to every month

## 3. Results

### 3.1 Descriptive analysis of the data

A total of 3,420 herds were registered with PRRS status during the study period, representing approximately 60% of the entire population. Herds are classified into several production types based on the ratio between sows and finishers (Schulz, 2019). Information on production type was missing for 277 herds. Among herds with known production types, finisher herds were the most common (69%), followed by sow herds (24%) and weaners (7%) (See Figure 3A). Among the study population, 1,804 herds were classified as SPF blue, 5 as SPF green, and 128 as SPF red, and 1,483 of non-SPF herds. Since SPF blue and SPF green herds follow the same detection process, they were treated identically in the model.

**Figure 3.**
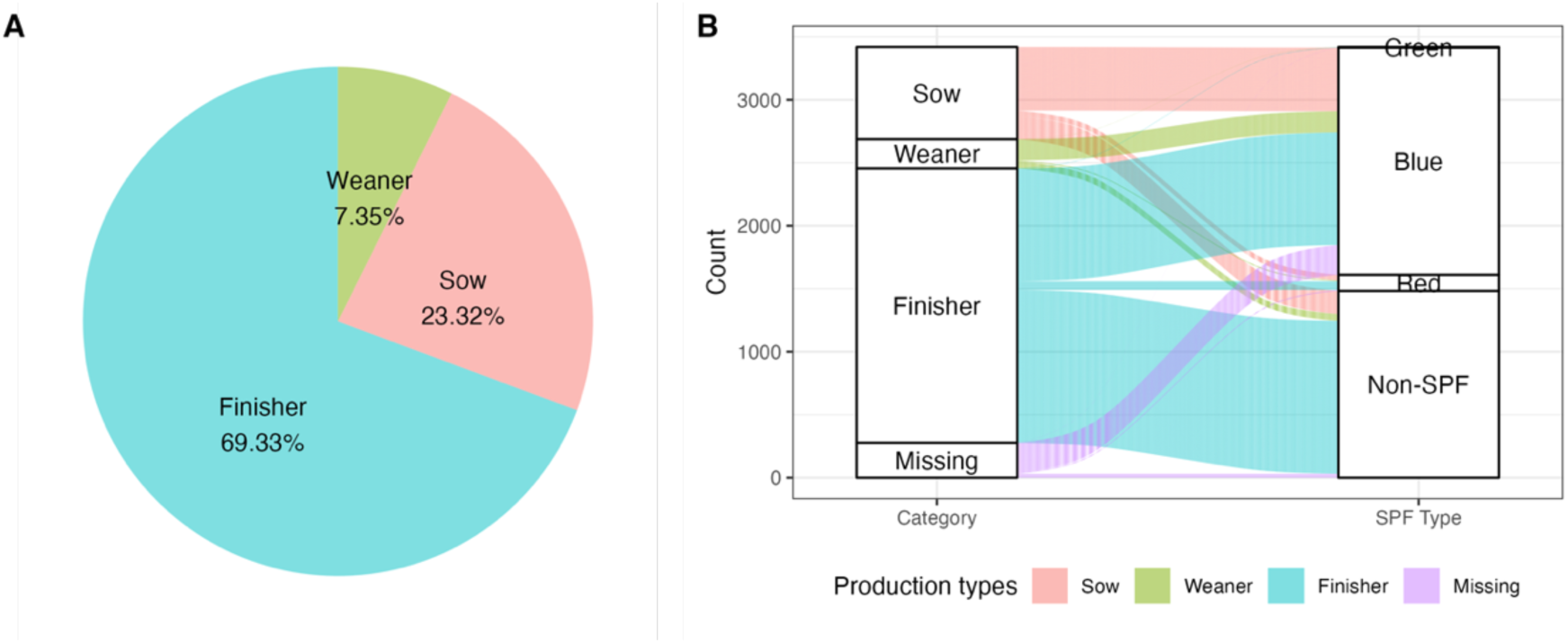
Composition of the study population: A) The percentage distribution of the three production types; B) the distribution of herds across production types and SPF status, showing the flow of herds into different biosecurity level (Green, blue, red, non-SPF). The thickness of the connecting lines represents the relative proportion of herds in each category.

The movement dataset consists of 100,464 records of movements, involving 1,682 sellers and 2,596 buyers between January 2020 and December 2021, with 15% (n = 498) of farms reporting no between-farm movement activities. Throughout the study period, 78 % of the pig movement relationships between pig farms persisted, and one fifth of the seller herds (n = 336) generated half of all pig movements. Most pig movements, among known production categories, occurred between sows and finisher facilities (34%) and between sows and weaner farms (15%). Overall, there was little variation in the distance traveled by pigs between farms across production types, with a median of 10.5 km (mean 35.6 km with a range 0–386.2 km). On average, herds have 11 neighboring herds in the local area (within 5km radius), with 25% percentile at 6 and 75% percentile at 16. Different production types show no significant differences in local density, though weaner herds are slightly more concentrated. Finisher herds have the fewest outward trading partners, averaging 2.26, while sow and weaner herds have more outward trading partners, averaging 2.57 and 3.14, respectively.

### 3.2 Between herd *R*_0_ map

Using the estimated parameters (Section 2.4), we calculate the between-herd basic reproduction ratio *R*_0_ and stratify it by the movement route *R*_*movement*_ and local spread *R*_*local* *spread*_. Herds with *R*_0_>1 are expected to transmit the infection to more than one other herd during their infectious period, while herds with *R*_0_ <1 are expected to transmit to fewer than one herd. The *R*_0_ map (Figure 4) shows that high risk herds are predominantly clustered in North Denmark, Central Denmark (especially along the coast), and certain areas of South Denmark. Our *R*_0_ map shows a strong alignment with observed prevalence. The distribution of between herd *R*_0_ is presented in Supplement Figure S5.

**Figure 4.**
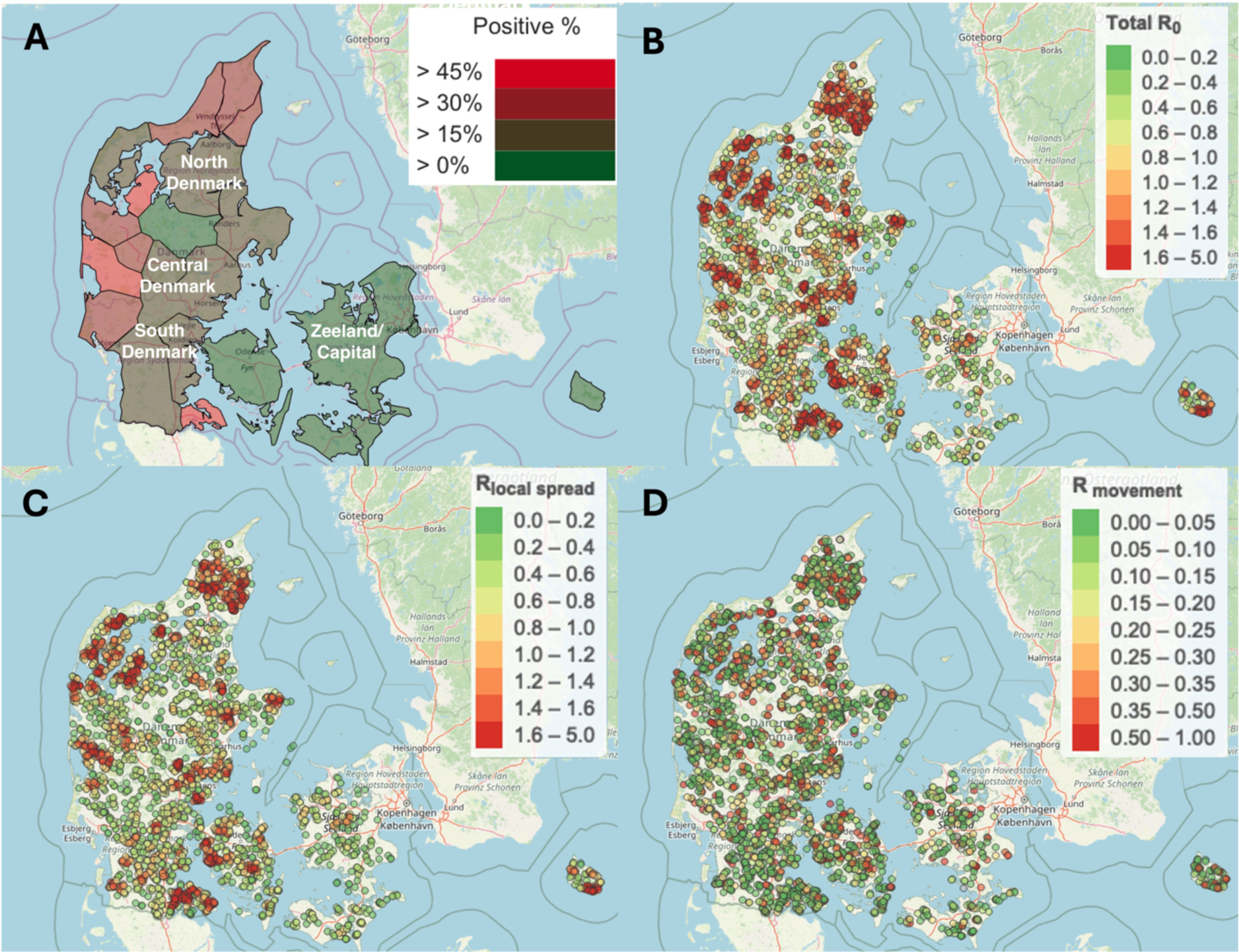
Between herd *R*_0_ map for the PRRS transmission. A) the observed prevalence in different regions in Denmark (Source from Landmand.dk); B) the between-herd R_0_ map; C) the *R*_*local* *spread*_map. Dots in red represent herds with R_0_ > 1 and dots in green and yellow represent herds with R_0_ < 1; D) *R*_*movment*_ map under 80% compliance with risk-based trading.

### 3.3 Relative contribution

Infectious herds progress through three phases, highly infectious (*I*_*h*_), lowly infectious phase (*I*_*l*_), and detected phase (*D*). In Section 2.5, we derive the between-herd *R*_0_ for each infectious phase, allowing us to express their relative contribution. For example, for the local spread, the three phases contribute to transmission with 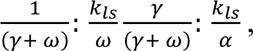 where herd production type and biosecurity level influence these ratios (Table 5). On average, detected herds contribute to around 40% of the total transmission. Parameters from Scenario 2 - 3 have only a marginal impact on this distribution, with contribution of detected herds ranging from 35% to 50%.

**Table 5.** Relative contributions of infectious phases, detection methods, and transmission routes across different production types and biosecurity levels using Scenario 1 and default herd sensitivity value. The “% of herds” column represents the proportion of herds for each production type and biosecurity level, summing to 100% across all categories. For each row, the relative contributions of *I*_*h*_(highly infectious), *I*_*l*_ (lowly infectious), and *D* (detected) sum to 100%. The "% detected by blood testing" column reflects the proportion of detected herds identified by blood testing, with the remainder detected via clinical notification (not shown in the table), together summing to 100%. The "% local spread" column represents the contribution of local transmission, with the remainder attributed to movements (not shown in the table), together summing to 100%.

| Production type | Biosecurity level | % of herds | % of $I_h$ contribution | % of $I_l$ contribution | % of $D$ contribution | % of local spread |
| --- | --- | --- | --- | --- | --- | --- |
| Sow | Red | 1% | 58% | 2% | 40% | 79% |
| Sow | Blue | 15% | 45% | 7% | 48% | 65% |
| Sow | Non-SPF | 5% | 33% | 8% | 59% | 70% |
| Weaner | Red | 0.2% | 59% | 4% | 37% | 65% |
| Weaner | Blue | 5% | 39% | 22% | 39% | 59% |
| Weaner | Non-SPF | 2% | 33% | 22% | 45% | 67% |
| Finisher | Red | 2% | 63% | 3% | 34% | 65% |
| Finisher | Blue | 33% | 41% | 24% | 35% | 63% |
| Finisher | Non-SPF | 37% | 37% | 24% | 39% | 70% |

In general, 59-79 % of the new infections are caused by the local spread, with an overall average contribution of 68%. The contribution tends to be highest in the sow herds, followed by the finisher herds and weaner herds. Transmission rate parameters can influence the relative contribution of the two routes: local spread contributes to approximately 70% of infections under Scenario 2, and 58% under Scenario 3.

The model includes both active surveillance (via blood sample testing) and passive surveillance (via clinical checkups). The proportion detected by blood sampling is given by competing rates of detection: 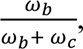 where two detection rates *ω* and *ω* are functions of the herd-level sensitivities of the clinical and blood-testing, as well as their frequency (See Eq. (5)). For red herds, which have monthly blood sampling, most of them are detected by blood testing (> 98%).

Using the herd-level sensitivities of clinical checkups from a recent study by Willkan et al. (2025), the model estimates that 35% of infected sow herds are detected by blood sampling and 66% by clinical checkups, while 79-88% of infections in weaner and finisher herds are detected by blood sampling with only 12%-21% detected by clinical checkups. However, if we match detection ratios similar to observed ratios from a report from SEGES innovation (2025), with approximately 70% of sow herd infections and 95% of weaner/finisher herd infections detected by blood testing, the herd level sensitivity for clinical checkups should be 3-4 fold lower (Table 2). However, the overall effect of herd-level sensitivity of clinical checkups on model output is limited, as shown in Supplement Figure S2.

### 3.4 Impact of Interventions

Observed herd prevalence stabilizes around 30% without additional interventions under the default control strategy in three scenarios (Figure 5). Despite variations in parameter assumptions, Scenarios 1-3 shows consistent trends in the impact of interventions.

**Figure 5.**
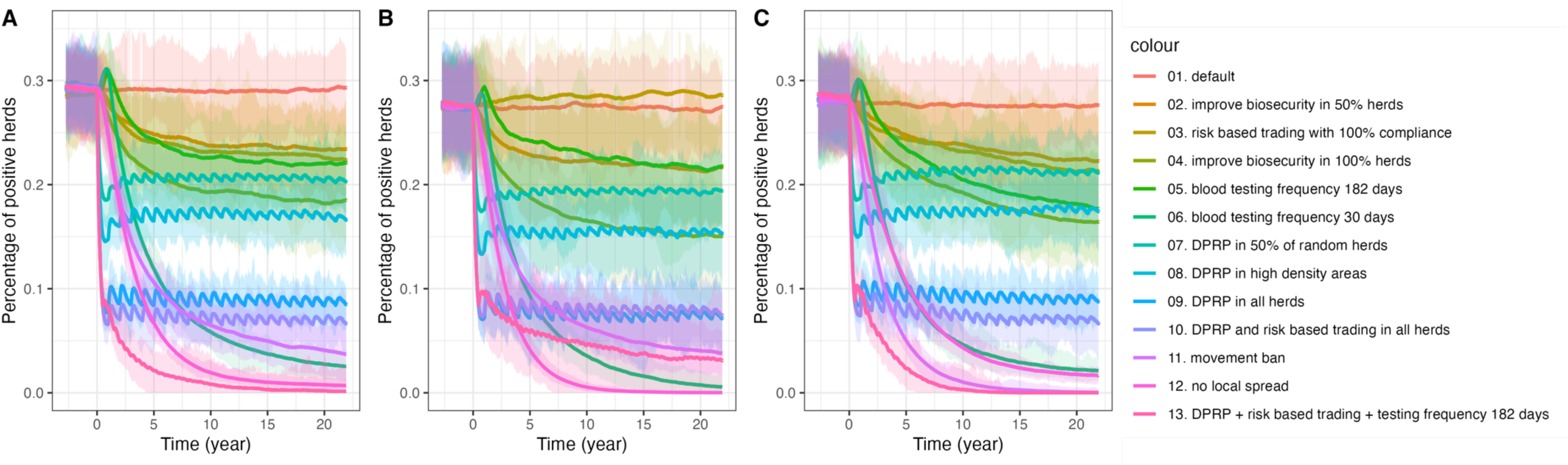
Impact of different intervention strategies on the observed prevalence across three scenarios: A) Scenario 1, B) Scenario 2, C) Scenario 3. Each line represents a specific intervention on combination of interventions, with colors corresponding to the legend on the right. The y-axis shows the percentage of positive herds, and the x-axis represents time in years. Shaded areas indicate variation across 100 model simulations. Interventions include biosecurity improvements, risk-based trading, depopulation-repopulation (DPRP) strategies, and combinations. All interventions are implemented starting from year 0.

With improving risk-based trading to full compliance, observed prevalence reduces to around 22% under Scenario 1 and 3 (Figure 5A&C, line 03), but increase to 28 % in Scenario 2. Risk-based trading cannot entirely prevent movement-mediated spread because of undetected infectious herds, as a hypothetic scenario complete movement ban can reduce prevalence to around 4% in Scenario 1 and 2 (Figure 5A & B, line 11) and 0.4% in Scenario 3 (Figure 5C, line 11).

Improving biosecurity by reducing herd susceptibility by 20% has only a modest impact with observed prevalence stabilizing at 23% when applied to half of the herds and 18% when implemented in all herds – consistently across three scenarios (Figure 5A-C, line 02 and 04). Increasing testing frequency to twice per year initially leads to a rise in observed prevalence and then reduce to around 20% (Figure 5A-C, line 05). Monthly testing has a much stronger effect, ultimately lowering prevalence to 2%-3% (Figure 5A-C, line 06).

Among all the interventions, depopulation-repopulation (DPRP) has the most immediate impact within the first two years, with prevalence declining to 10%-20% (Figure 5, line 07-09), depending on the proportion of herds complying with the measure. The combination of depopulation-repopulation, stricter risk-based trading and improving blood testing frequency to twice per year proves to be sufficient in eradicating PRRS within 20 years on average (Figure 5A & C, line 13), but not in Scenario 2 (Figure 5B, line 13). These results highlight that while individual interventions can reduce transmission to some extent, a comprehensive approach might be needed for eradication.

Using the simulated true prevalence (see Supplement Figure S6) and the equation in Section 2.5, the national-level basic reproduction number *R*_0_ are estimated (Table 6). Whether PRRS can persist for each control strategy depends on whether *R*_0_ is below 1. Either local spread or movement alone may be sufficient to sustain PRRS transmission, as indicated by *R*_0_> 1 under control strategies 11 and 12 (Table 6).

**Table 6.**
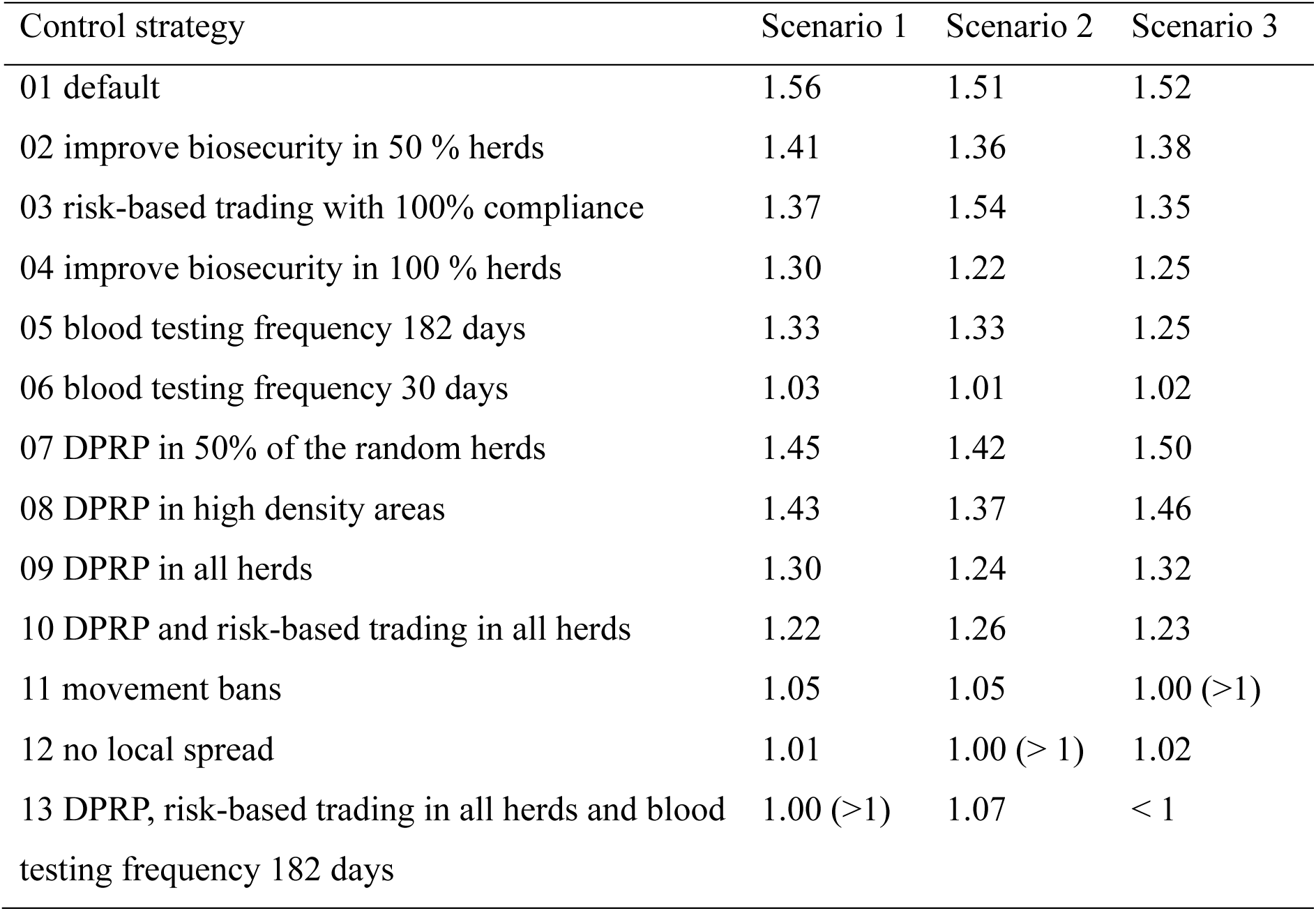
National *R*_0_ under different control strategies and scenarios.

Controlling local spread via depopulation-repopulation (DPRP) might seem effective in reducing the observed prevalence (Figure 5 line 09), but its impact on the true prevalence and *R*_0_are limited (Table 6 line 09). In comparison, increasing the frequency of blood testing frequency has the strongest effect in reducing transmission (Table 6 line 06), even though it can initially lead to an increase in observed prevalence due to improved detection (Figure 5 line 06).

## 4. Discussion

PRRS has been endemic in Denmark for over 30 years, causing significant economic losses, with the estimated cost ranging between 4 to 139 euro per sow per year (Rathkjen & Dall, 2017). In 2022, a reduction plan was introduced, focusing on compulsory PRRS-status reporting, elimination of PRRS in positive herds and recommended risk-based trading under SPF system. This raises questions about the effectiveness of these measures. To address this, we developed a between-herd transmission model accounting for both local spread and movement-mediated transmission, replicating dynamics before the legislative change and predicting control impacts.

A key innovation of this study is its modeling of PRRS spread, considering the distinct roles of highly infectious and lowly positive herds and explicitly incorporating the Danish surveillance system. The model estimated that highly infectious herds (positive unstable) are 50 times more infectious than lowly infectious herds (positive stable), highlighting their important role in PRRS transmission and control. In addition, we assessed that the current surveillance program misses 16% of infectious herds, which may contribute up to 60% of the overall transmission. These undetected infections can be reduced by more frequent testing, aligning with recent findings on benefits of increased sampling frequency for early detection (Willkan et al., 2025). Since herds are most infectious during the first three months after infection, testing at intervals of three months or shorter can substantially reduce the number of hidden infections, as illustrated by the comparison between lines 10 and 13 in Figure 5.

This study generates the first between-herd *R*_0_ map for the PRRS, explaining the spatial heterogeneity in PRRS transmission. The findings are consistent with observed high prevalence in certain Danish regions including North Demark, Central Denmark (especially along the coast), and parts of South Denmark (Figure 4). A previous study also identified significant spatial clusters, suggesting PRRS endemic states in these areas (Lopes Antunes et al., 2015). Stratification of *R*_0_reveals distinct spatial pattern: *R*_*mov*_is relatively homogenous across the country, whereas *R*_*local* *spread*_shows strong spatial clustering, reflecting spatial patterns in observed surveillance data (Figure 4). This suggests that local spread is a key driver of PRRS transmission in Denmark. This is further supported by the estimates of approximately 60-70% of new infections caused by local spread (Table 5). Although scenarios 2 and 3 increase the relative contribution of movement-related transmission to 40%-50%, local spread remains a dominant role. However, it is important to note that our study includes all the herds that report their PRRS status, representing 60% of the total population. Consequently, some local spread may in fact reflect movements involving those non-included herds. With the eradication plan requiring compulsory PRRS status reporting for herds with more than 10 sows or 100 pigs, future analyses should include a broader range of herds when available and could help clarify this uncertainty and improve our understanding.

Several risk factor analyses have identified high density as a main risk factor, suggesting that local spread – potentially through airborne transmission– plays an important role (Arruda et al., 2017; Baysinger et al., 1997; Firkins & Weigel, 2004; Mortensen et al., 2002). Modelling studies in the U.S. have estimated that pig movement explained 15 to 20% of the new infections, while 50%-60% due to local spread in sow and finisher herds (Galvis, Corzo, & Machado, 2022; Galvis, Corzo, Prada, et al., 2022). These studies also found that pig movements played a greater role in weaner herds but were less in finisher and sow herds. The pattern aligns with our modeling results. Approximately 70% of Danish herds are finishers, and since they primarily sell pigs to slaughterhouses, their role in movement-mediated transmission is limited. Compared to these studies, our model simplifies indirect contact as part of local spread due to data limitations. However, this simplification is unlikely to have a significant impact, as indirect contact accounts for only 0.3% to 2.5% of transmission (Galvis, Corzo, & Machado, 2022). In contrast, studies in Canada have attributed PRRS transmission primarily to direct or indirect contact between herds under shared ownership as they found no evidence of airborne transmission (Rosendal et al., 2014). As a result, modeling efforts in Canada have focused solely on movement-mediated transmission (Thakur et al., 2015). These cross-country differences may be explained by variations in herd density, herd structure, and control measures, such as risk-based trading in Denmark. Despite multiple studies highlighting the role of animal movement in PRRS spread (Amirpour Haredasht et al., 2017; Pamornchainavakul et al., 2023; Thakur, Sanchez, et al., 2015), our model is the first to assess the impact of risk-based trading. We showed that with 80% of compliance, risk-based trading reduced the observed prevalence from 40% to 30% and can further reduce it to 20% with 100% of compliance.

High density of herds, complex trade patterns, infrequent active surveillance and low sensitivity of passive surveillance all contribute to the persistence of PRRS (as shown in Table 5). Depopulation-repopulation shows an immediate and strong effect on observed prevalence, but its impact on true underlying transmission is limited as it does not influence hidden infections. In contrast, increasing testing frequency has the greatest effect on the reducing the real transmission, although it initially raises observed prevalence. Our results suggest that interventions targeting different transmission routes can have an additive effect. A comprehensive control should include measures to reduce hidden infections via more frequent testing, control the local spread in high-density areas via improving biosecurity measures, depopulation-repopulation, stricter risk-based trading to limit the introduction of PRRS (Figure 5, line 03) in free areas (Figure 5, line 05 and 13). Similar suggestions have been made in Germany (Fahrion et al., 2014).

An overestimation of *R*_0_ and an underestimation of intervention effectiveness in the present study could be caused by the higher proportion of false-positive test results in the current surveillance system. Specifically, herds that purchase pigs from PRRS-positive herds are classified as PRRS-positive, introducing uncertainty in estimating the true infection probability from movement. False-positive herds in the surveillance data do not contribute to further PRRS transmission, whereas assuming an infection probability of 1 artificially increases the transmission potential. To assess the impact of assuming a 100% infection probability via risky movement for both highly infectious and lowly infectious herds, we tested this scenario, which led to an overestimation of PRRS prevalence at around 43%, compared to the observed prevalence of 30%. This suggests that the infection probability via risky movement is likely lower than 1.

Further, a characteristic of the Danish pig industry is the establishment of joint operations to reduce testing costs and mitigate transmission risks (SPF, 2024). Under this system, multiple herds submit samples together, and their PRRS status is updated simultaneously in the surveillance system. Herds within a joint operation can be automatically classified as positive if any other herd in the group tests positive. A similar issue exists in the Danish cattle industry for Salmonella surveillance, where classifications are updated at the business level rather than for individual farms (Conrady et al., 2024). Pig herds within a joint operation typically share the same owner, engage in regular trade (e.g., from sow herds to weaner herds and then to finisher herds), or are located in close proximity. Given the higher likelihood of transmission between these connected herds, it is reasonable to treat them as a single epidemiological unit. Despite functioning as a unified unit epidemiologically, each herd within the joint operation still maintains individual entries in the surveillance system. This leads to an overestimation of the observed prevalence. Relying on this biased surveillance data to assess transmission dynamics can result in an overestimation of *R*_0_, making PRRS appear more difficult to eradicate in the model than it is.

This model was calibrated with PRRS-1 surveillance data, as PRRS-1 is more prevalent in Denmark (Strategy for the Reduction of PRRS in Pigs in Denmark, 2022), while PRRS-2 can be associated with the vaccination strain was originally introduced with a vaccination program (Balka et al., 2015; Kvisgaard, 2017). Experimental studies suggest that vaccination can protect animals from re-infection (Canelli, 2018; Lunney et al., 2010), but its efficacy in preventing transmission at herd level remains unclear and controversial. Vaccine strains have been shown to spread between farms (Bøtner et al., 1997, 1999; Grosse Beilage et al., 2009; Kiss et al., 2006), raising concerns about their role in transmission. Modelling studies assumed 1% of vaccine efficacy at the herd level, leading to only a marginal decrease in incidence (Galvis, Corzo, Prada, et al., 2022; Valdes-Donoso & Jarvis, 2022). In Denmark, modified live viral vaccines for PRRS-1 and PRRS-2 are used exclusively in PRRS-positive herds to mitigate clinical symptoms and are not considered a tool for reducing between herd transmission. Therefore, we did not model the impact of vaccination.

While our model sufficiently captures spatial heterogeneity at a national scale, we acknowledge that the model and data have limitations in representing all aspect of spatial heterogeneity. As a herd-level transmission model, it cannot assess the impact of herd size or internal biosecurity measures on the within-herd transmission dynamics, such as the number of pigs getting infectious or the duration of outbreak in a herd (Evans et al., 2010). In addition, while the model explicitly accounts for herd owners’ decisions on depopulation-repopulation and improving biosecurity level, the lack of data at herd level requires assuming these decisions. Although SPF herds in Denmark are recommended to quarantine incoming gilts for 42 days—potentially reducing infection risk via movements - there is no available data to quantify transmission probability through risky movements (*P*_*mov*_). As a result, we are unable to assess the effectiveness of quarantine in reducing PRRS transmission with the current surveillance data. Future studies that monitor herd status changes after risky movement, both with and without quarantine, could improve our understanding of how quarantine impact the control of PRRS. Nonetheless, our study suggests that achieving eradication may require a combination of more frequent testing, targeted within-herd PRRS elimination, and stricter risk-based trading practices. It also identifies PRRS hotspots and transmission routes, providing evidence-based recommendations for effective control.

## Conclusion and recommendation

This study identifies high-risk areas for PRRS transmission presented in *R*_0_ map. In the study population, current surveillance misses about 17% of the infectious herds that contribute to 60% of the transmission. While PRRS elimination within herds via depopulation-repopulation seems the most effective control measure on reducing the observed prevalence, it might have limited impact on reducing the true transmission, as large part of the transmission is attributed to hidden infection. A comprehensive control strategy is essential for nationwide eradication. Improved active surveillance through more frequent testing, prioritizing local control in high-risk areas, stricter risk-based trading to protect PRRS-free areas are key to achieving eradication.

## Acknowledgement

We would like to thank Stefan Widgren, Pauline Ezanno, Nicolai Rosager Weber, Sara Dalsgaard, and Kristian Møller for their invaluable suggestions contributed to the improvement of this study. This work received co-funding from European Union’s Horizon Europe under Grant Agreement No 101136346: European Partnership Animal Health &Welfare, Joint Internal Project SOA10, SOA 22 and SOA 15 and the pigs levy fund.

## Supplementary Materials

### S1. Illustration of the SPF system movement restriction

**Figure S1.**
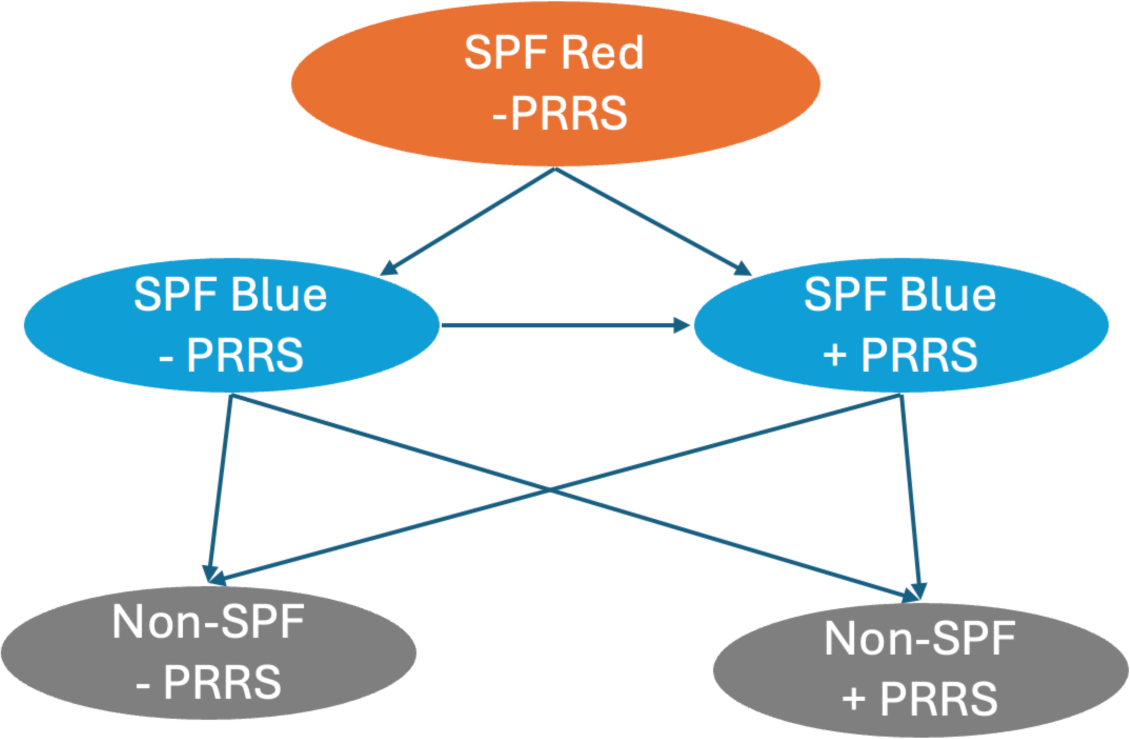
Illustration of the movement restriction in the SPF system. Herd owners can choose to follow this scheme to maintain their PRRS status. However, purchasing pigs from herds with a lower status will result in losing their PRRS status.

### S2. Sensitivity analysis for the herd sensitivity of the clinical checkups

**Figure S2.**
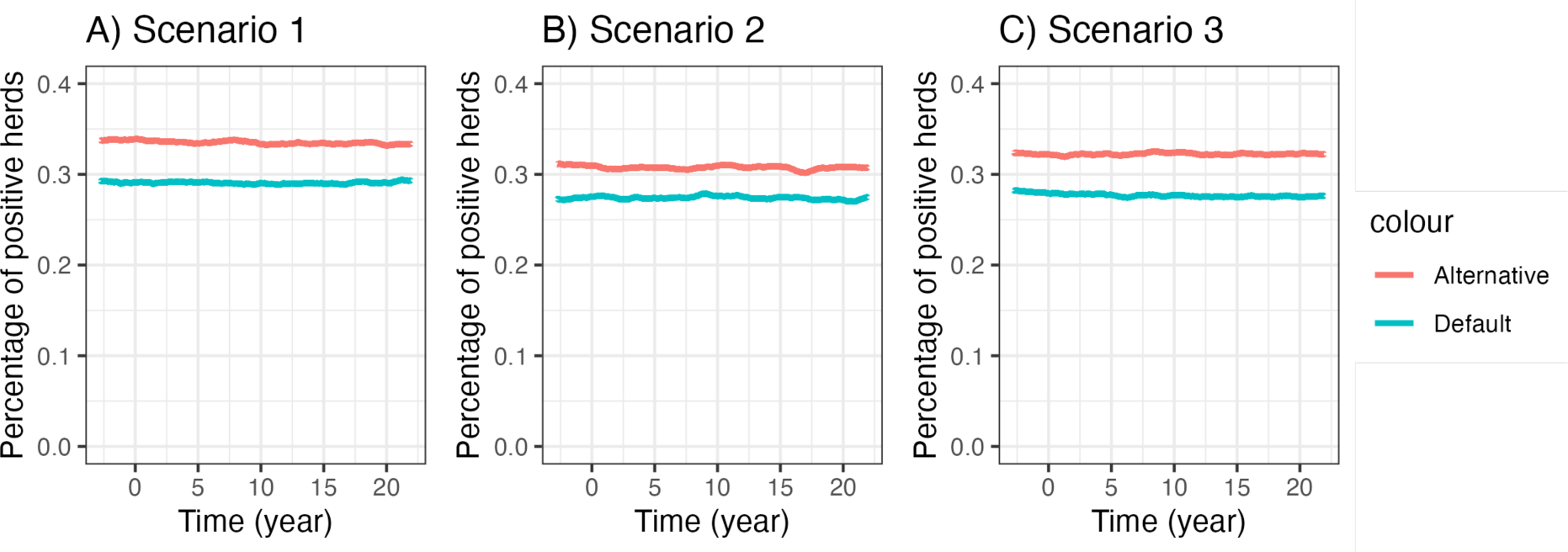
Sensitivity analysis of herd-level sensitivity for clinical checkups. The default herd sensitivity is 0.30 for sow herds and 0.05 for weaner/finisher herds (Willkan et al., 2025). In the alternative scenario, herd sensitivity is reduced to 0.07 for sow herds and 0.015 for weaner/finisher herds (SEGES Innovation, 2025). The lower sensitivity values result in a slightly higher proportion of positive herds detected compared to the default setting.

### S3. Confidence interval of estimated transmission rate parameters

**Table S1.** Estimation of transmission rate parameter for local spread.

| Parameter | Definition | Point estimation | 95% confidence interval bound |
| --- | --- | --- | --- |
| $\beta_{ls}$ | Local spread transmission rate parameter for highly infectious herds | 1.58e-3 | (3.1e-4, 3.0e-3) |
| $K_{ls}\beta_{ls}$ | Local spread transmission rate parameter for lowly infectious herds | 1.74e-5 | (2.53e-4, 3.67e-4) |

### S4. Calculation of *R*_0_

For each transmission route, the basic reproduction ratio can be calculated using the transmission matrix, using *R*_*local* *spread*_ as an example.

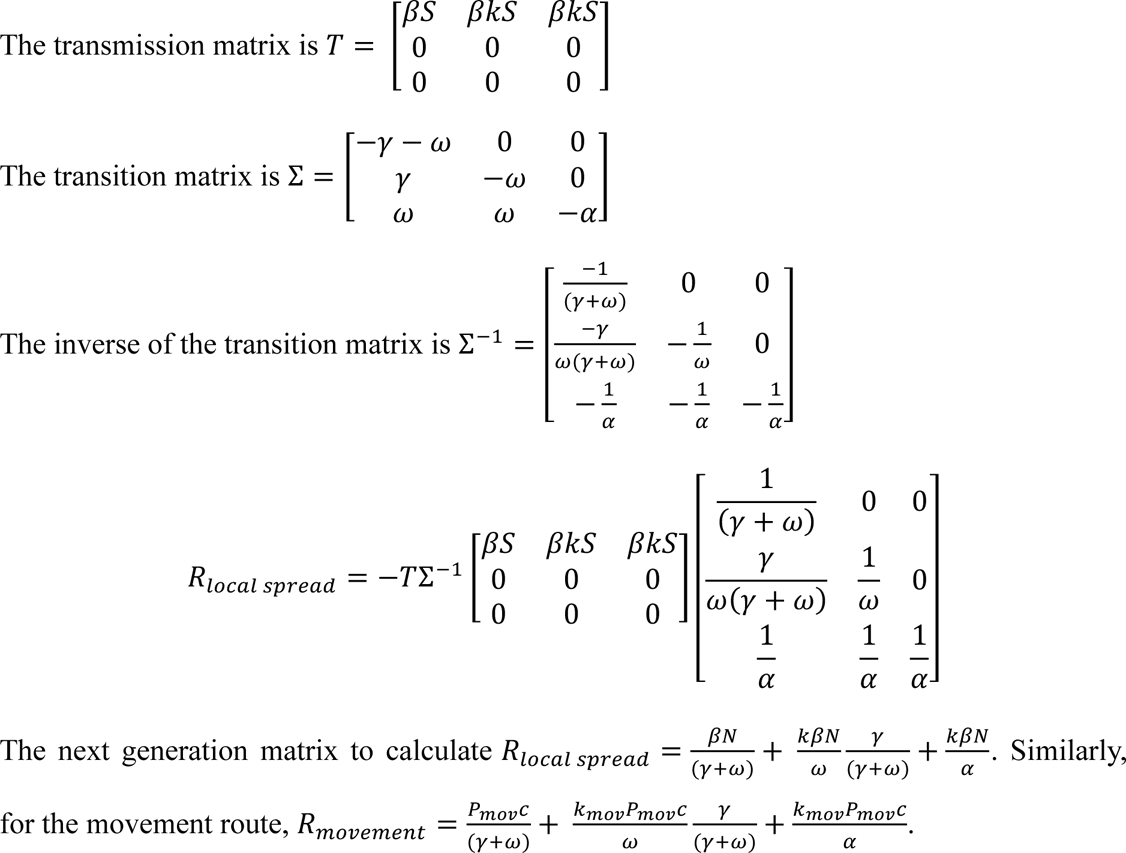

### S5. Between herd *R*_0_ distribution

**Figure S5.**
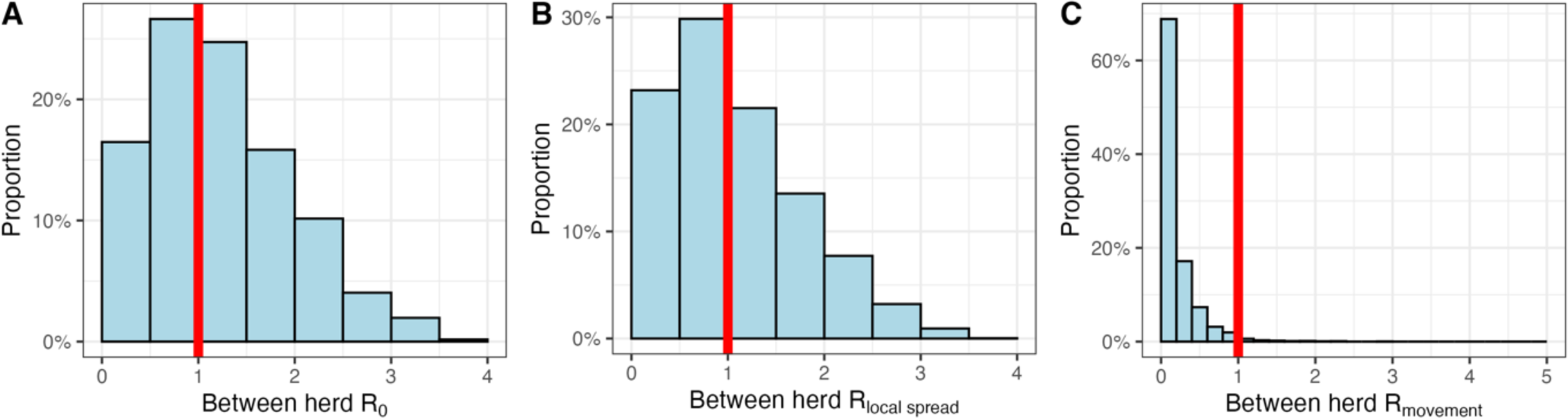
Distribution of the between-herd basic reproduction ratio *R*_0_ for all herds in the study population. A) total between-herd *R*_0_; B) *R*_*local* *spread*_; C) *R*_*movement*_

In the study population, 57% of herds have a between-herd reproduction ratio *R*_0_ greater than 1. Local spread alone results in 47% of herds infecting more than one herds (*R*_*local* *spread*_ > 1). In comparison, fewer than 0.2% of herds have *R*_*movment*_>1, as the average number of trading partners per herd is below 1 under current risk-based trading compliance.

### S6. Simulated true prevalence under three scenarios

**Figure S6.**
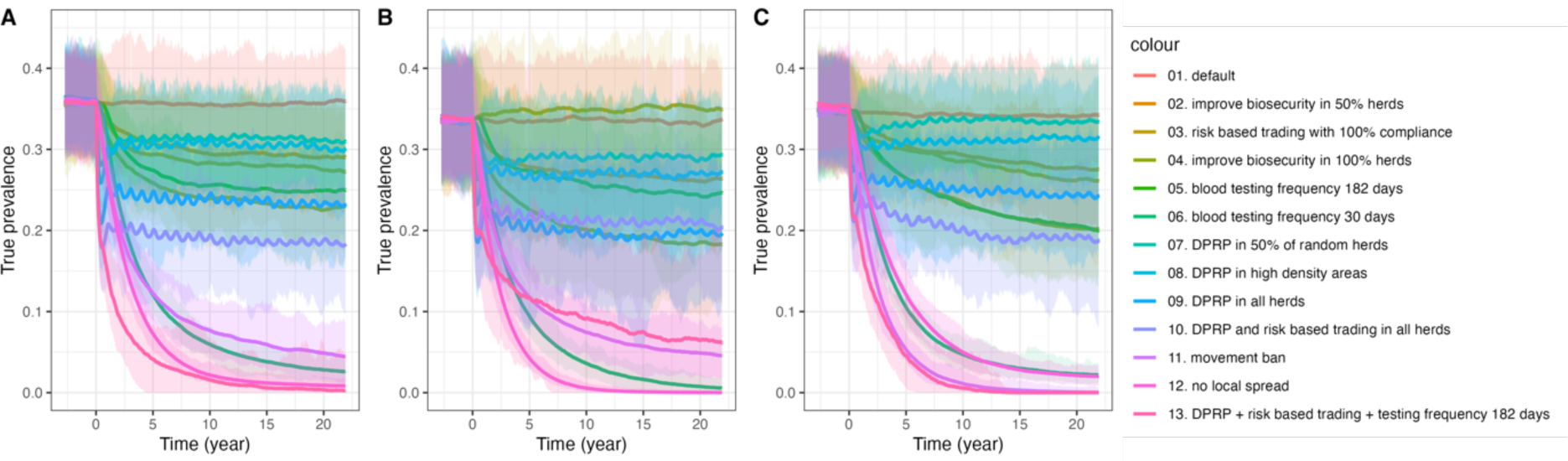
Impact of different intervention strategies on the true prevalence across three scenarios: A) Scenario 1, B) of Scenario 2, C) Scenario 3. Each line represents a specific intervention on combination of interventions, with colors corresponding to the legend on the right. The y-axis shows the percentage of positive herds, and the x-axis represents time in years. Shaded areas indicate variation across 100 model simulations. Interventions include biosecurity improvements, risk-based trading, depopulation-repopulation (DPRP) strategies, and combinations. All interventions are implemented starting from year 0.

## Notes

### Competing Interest Statement

The authors have declared no competing interest.

### Summary of Updates

Typo in highlight and author names

